# Spatial coupling of endogenous Tau translation and degradation by neuroproteasomes in dendrites revealed by STARFISH

**DOI:** 10.1101/2025.06.04.657939

**Authors:** KD Konrad-Vicario, V Paradise, L Demir, CD Makinson, Z Ming, KV Ramachandran

**Affiliations:** Taub Institute for Research on Alzheimer’s Disease and the Aging Brain, Columbia University, New York, NY 10032; Department of Neurology, Vagelos College of Physicians and Surgeons, Columbia University, New York, NY 10032; Department of Neuroscience, Vagelos College of Physicians and Surgeons, Columbia University, New York, NY 10032; Synbio Technologies, Monmouth Junction, NJ 08852

## Abstract

Cells regulate protein synthesis, folding, and degradation to maintain proteostasis, and disruptions in these processes have been linked to neurodegenerative diseases. In Alzheimer’s disease (AD), the protein Tau mislocalizes from axons to the somatodendritic compartment and aggregates into pathological filaments. Although Tau aggregation is a hallmark of AD, the subcellular dynamics of its synthesis and degradation are not well characterized. Because nascent polypeptides are particularly susceptible to misfolding, local control of Tau synthesis and degradation may be essential to prevent aggregation. Here, we develop STARFISH, a method for visualizing the subcellular site of endogenous mRNA translation in primary neurons and *in vivo* with single-molecule sensitivity and near-codon resolution, without modifying the nascent polypeptide. Using STARFISH, we show that despite the broad distribution of *Mapt* mRNA, Tau is translated almost exclusively in neuronal dendrites, revealing an unexpected level of spatial regulation. We further identify that one-third of newly synthesized Tau is co- or peri-translationally degraded in dendrites by a neuronal-specific plasma membrane-associated proteasome, the neuroproteasome. Failure of neuroproteasome-mediated degradation leads to the protein synthesis-dependent accumulation of somatodendritically mislocalized Tau aggregates. These findings define a previously unrecognized proteostasis mechanism that counterbalances the constitutive physiological overproduction of Tau. We speculate that failure of this proteostasis system contributes directly to Tau aggregation in dendrites, defining a new pathomechanism in Alzheimer’s disease.

## Introduction

Protein translation is highly tuned to cellular demands. Beyond simple modulation of protein levels, translation of specific mRNAs can be spatially controlled^1^. Spatially compartmentalized translation is exemplified in neurons, where protein synthesis occurs in distal locations, away from the soma, in order to respond to local demands such as synaptic activity^2^. However, nascent polypeptides, either during or immediately following translation, are metastable and are particularly susceptible to misfolding and aggregation^3^. Rapid and compartmentalized protein synthesis in neurons therefore presents a significant risk for proteostasis collapse and protein aggregation in distal processes. The mechanisms that act as counterbalances to local and rapid protein synthesis demand remain obscure.

Many neurodegenerative diseases are characterized by specific protein aggregates. It is unclear whether aberrant proteostasis of specific disease-relevant nascent polypeptides contributes to their aggregation. We recently reported that failure of a neuronal-specific proteostasis mechanism, a plasma-membrane associated proteasome we term the neuroproteasome, results in the formation of endogenous Tau paired helical filaments (Paradise, Konrad-Vicario et al, In Revision). In a previous report, we provided evidence that neuroproteasomes degrade newly synthesized proteins^4^. Taken together, we hypothesized that neuroproteasomes degrade newly synthesized Tau and this failure results in AD-relevant Tau aggregates.

A unique feature of Tau deposition in the AD brain is strong enrichment and mislocalization of hyperphosphorylated Tau in the soma and dendrites, compared to the enrichment of Tau in axons in the physiological state^5-9^. The conventional explanation for this observation is that axonal Tau becomes phosphorylated, detaches from the microtubule, and translocates into the somatodendritic compartment^5,10^. Multiple groups have shown that Tau protein is present postsynaptically and that Aβ oligomers, as well as neuronal activity, result in a protein synthesis-dependent elevation of Tau levels in the somatodendritic compartment^9,11-13^. However, the dynamics of Tau synthesis, and their relevance to Tau aggregation, remain vague.

One impediment to this analysis has been a high-resolution tool to visualize translation of endogenous mRNAs. Many tools exist to visualize protein synthesis in cells, but the majority require either modifying the nascent polypeptide with puromycin or unnatural amino acids or using reporter mRNAs to visualize active translation^14^. Modification of mRNAs or proteins can change their subcellular distribution and many methods do not visualize the endogenous transcript^15-17^. Furthermore, puromycin treatment evicts translating ribosomes off the mRNA and the puromycylated peptide can diffuse, making it challenging to visualize the site of protein synthesis with codon-level resolution^18-20^. These puromycylated truncated products can also act as DRiPs, which impact the measured turnover kinetics^21,22^. Finally, new methods to visualize the subcellular site of translation require denaturation or precipitation of proteins, and therefore are not compatible with labeled cells or subsequent antibody labeling.

## Results

### STARFISH reveals the subcellular localization of translation with near-codon resolution

We recognized a need to develop high resolution methods to visualize with high spatial precision where an mRNA was being translated, without modifying the nascent polypeptide or mRNA sequence. We therefore coupled single-molecule FISH experiments with ultra-efficient proximity ligation assays (PLA)^23^ between the 18S rRNA and a target mRNA^24^ using clickable probes to increase reliability and sensitivity (**Fig. 1a**). We carefully designed the probes to minimize the distance required between probes to provide a positive PLA signal, in an effort to improve spatial precision and visualize ribosomes as close to the decoding center as possible. We refer to this technique to visualize ribosomes bound to mRNAs as <u>s</u>patially-resolved tr<u>a</u>nslation of endogenous <u>R</u>NA FISH, or STARFISH.

**Figure 1:**
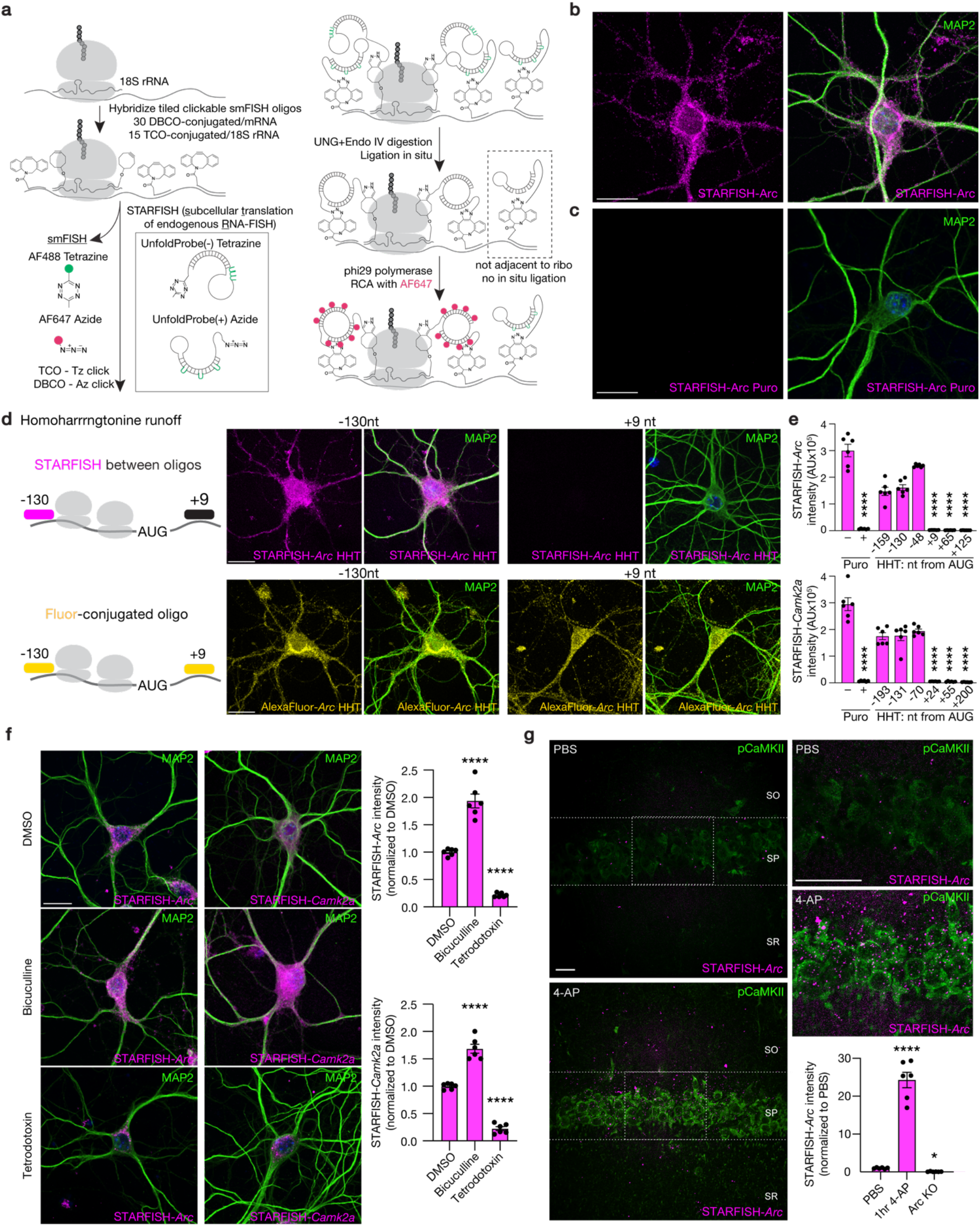
STARFISH measures the subcellular site of translation with near-codon resolution. **a**, Schematic of STARFISH (subcellular translation of endogenous RNA-FISH) experiments to visualize subcellular site of mRNA translation. **b**, STARFISH against *Arc* mRNA (encoding Arc) in DIV18 primary hippocampal neurons. STARFISH-*Arc* signal (Magenta), MAP2 (Green), DAPI (Blue). **c**, STARFISH-*Arc* signal after Puromycin (Puro) pulse, which dissociates translating ribosomes off mRNAs. **d**, Homoharringtonine (HHT) runoff experiments, schematic to left indicates ribosomes in HHT-treated neurons at initiating AUG unable to undergo a new round of translation. STARFISH (Magenta) compared to Fluor-conjugated probes (Yellow), DAPI (blue) in merge. Single mRNA probes used for all experiments; mRNA probe position relative to the initiating AUG indicated. **e**, Quantification of STARFISH-*Arc* and STARFISH-*Camk2a* intensities (Extended Data Fig 1b). Puromycin (Puro) and HHT experiments indicated, tiled probes used for Puro, single probes used for HHT bound to indicated positions relative to AUG. N=6 biological replicates. **f**, STARFISH-*Arc* and STARFISH-*Camk2a* following stimulation of neuronal activity with Bicuculline (Bic, 1*μ*M, 15 min) or suppression with Tetrodotoxin (TTX, 1*μ*M, 15 min). Quantification of normalized STARFISH intensity signals relative to DMSO to right. N=6 biological replicates. **g**, STARFISH-Arc in hippocampi from mice intraperitoneally injected with 4-AP compared to PBS control. Quantification of STARFISH-*Arc* signal normalized to PBS plotted to right. N=6 biological replicates. All scale bars are 20 µm. **e**,**f**,**g** ****p<0.001 by One-Way ANOVA Tukey’s Multiple Comparison Test,

As a first test, we measured STARFISH against Arc, which has been demonstrated to be locally translated in neurons. We observe broad somatic and dendritic STARFISH-*Arc* (Arc) signal in primary mouse hippocampal neurons with single molecule sensitivity (**Fig. 1b, Extended Data 1a**). Upon addition of puromycin, we see a complete loss of STARFISH signal, as expected for ribosome-bound mRNAs (**Fig. 1c**). We then performed STARFISH against two other mRNAs, *Camk2a* and *Actb*. We detect strong puromycin-sensitive signal for both STARFISH-*Camk2a* and STARFISH-*Actb* throughout neuronal processes and in soma (**Extended Data Fig. 1b-e**).

In order to measure the maximal proximity between a ribosome and the mRNA probe necessary to produce a STARFISH signal, we exploited the property that Homoharringtonine (HHT) leads to stalling of ribosomes precisely at the initiating AUG^25^. In HHT ribosome runoff assays, we used probes binding to a single segment of the mRNA sequence instead of multiple probes tiling the mRNA sequence in order to precisely define the distance between Ribosome and mRNA probe required to generate a STARFISH signal. We used probes as close as possible to the initiating AUG without losing specificity for target mRNAs. After HHT treatment, we fail to detect any STARFISH signal using mRNA probes 9 nucleotides or farther from the initiating AUG, while probes in the 5’ UTR still provide robust signal (**Fig. 1d,e, Extended Data Fig 2a**). HHT runoff assays eliminate the STARFISH signal in the CDS, but do not affect the ability of the probes to bind (**Fig 1d,e, Extended Data Fig 2a**), demonstrating that STARFISH likely only measures intramolecular interactions rather than intermolecular interactions. We find similar resolution for other tested mRNAs, such as *Camk2a* (**Extended Data Fig 2b**).

The signal upon HHT or Puromycin treatment is nearly undetectable: STARFISH generates over 100-fold greater signal relative to puromycin or HHT controls (**Fig. 1e, Extended Data Fig 2**). In comparison, current methods give approximately 2-3 fold signal relative to these controls and the puromycin and HHT controls do not fully eliminate signal, which suggests that some of the signal from these methods comes from non-translating ribosomes^24,26^. Our data support the conclusion that STARFISH measures the localization of actively translating ribosomes of endogenous mRNAs with higher signal/noise and higher spatial resolution compared to existing methods (∼3 codon relative to 40+). Since STARFISH only requires paraformaldehyde fixation and not methanol, STARFISH maintains protein epitopes and is compatible with post hoc identification of labelled cells and immunocytochemistry.

### STARFISH detects activity-dependent translation in primary neurons and *in vivo*

It is well established that certain proteins are translated in neurons in response to neuronal activity. To validate the ability of STARFISH to visualize activity-dependent translation, we measured STARFISH against the *Arc* and *Camk2a* mRNAs, for which there is strong evidence of activity-dependent translation^27-30^. We found a significant suppression of basal STARFISH-*Arc* and STARFISH-*Camk2a* signals after inhibition of sodium channels with Tetrodotoxin (TTX) in primary hippocampal cultures (**Fig. 1f**). Conversely, we observed a robust increase in STARFISH-*Arc* and STARFISH-*Camk2a* signal minutes after bicuculline addition to primary hippocampal cultures (**Fig. 1f**). Modifying neuronal activity had no effect on STARFISH-*Actb* (**Extended Data Fig. 3a**).

We next tested whether we could detect activity-dependent translation in *ex vivo* slice preparations and *in vivo*. We stimulated Schaffer collaterals to induce robust activity in CA1 pyramidal neurons, a paradigm which induces an activity-dependent increase in CamKII and Arc mRNA and protein levels^27,31^. We observed a significant increase in STARFISH-*Arc* and - *Camk2a* signals in CA1 immediately after Theta Burst Stimulation, which is absent in slices incubated with aniosomycin (**Extended Data Fig. 3b**). We do not detect an elevation of STARFISH-Actb in these experiments (**Extended Data Fig. 3b**). Finally, to test the ability of STARFISH to detect translation *in vivo*, we augmented neuronal activity *in vivo* by intraperitoneal injections of low-dose 4-aminpopyridine (4-AP)^32^. We again observe intense STARFISH-*Arc* staining in brains of animals injected with 4-AP compared to vehicle injected controls (**Fig. 1g**). Together, our data support that STARFISH is able to visualize relative changes in translation in response to neuronal activity.

### Tau is synthesized in neuronal dendrites

We next sought to reveal examples of where STARFISH could gain valuable biological insights beyond activity-triggered translation. There are many discrepancies between mRNA localization and protein localization^33^. For example, Tau is an axonally localized protein, but using single molecule FISH experiments, it is widely reported that *Mapt* mRNA (the mRNA encoding for Tau) is distributed throughout the somatodendritic compartment^34,35^. Indeed, some groups have reported somatodendritic distribution of Tau protein, but this has been highly dependent on the antibody used and the phosphorylation status of Tau^9,12,13,36^. Given the resolution and sensitivity of STARFISH, we considered STARFISH against *Mapt* may shed light into the discrepancy between mRNA and protein localization.

First, we developed probes tiling the *Mapt* transcript and performed single-molecule FISH experiments to determine the localization of *Mapt* mRNA. Consistent with previous observations, we find *Mapt* mRNA throughout the neuron, including the somatodendritic compartment (**Fig. 2a**) compared to secondary-only and Tau-KO controls (**Extended Data Fig. 4a, b**). Using the same probes, STARFISH analyses reveal that Tau translation occurs within the dendrite, with no reliable signal in the soma or in axons (**Fig. 2b, Extended Data Fig. 4c**), despite the abundance of both *Mapt* mRNA and ribosomes in the soma. STARFISH-*Mapt* is sensitive to puromycin and HHT-runoff controls (**Extended Data Fig. 4d**).

**Figure 2:**
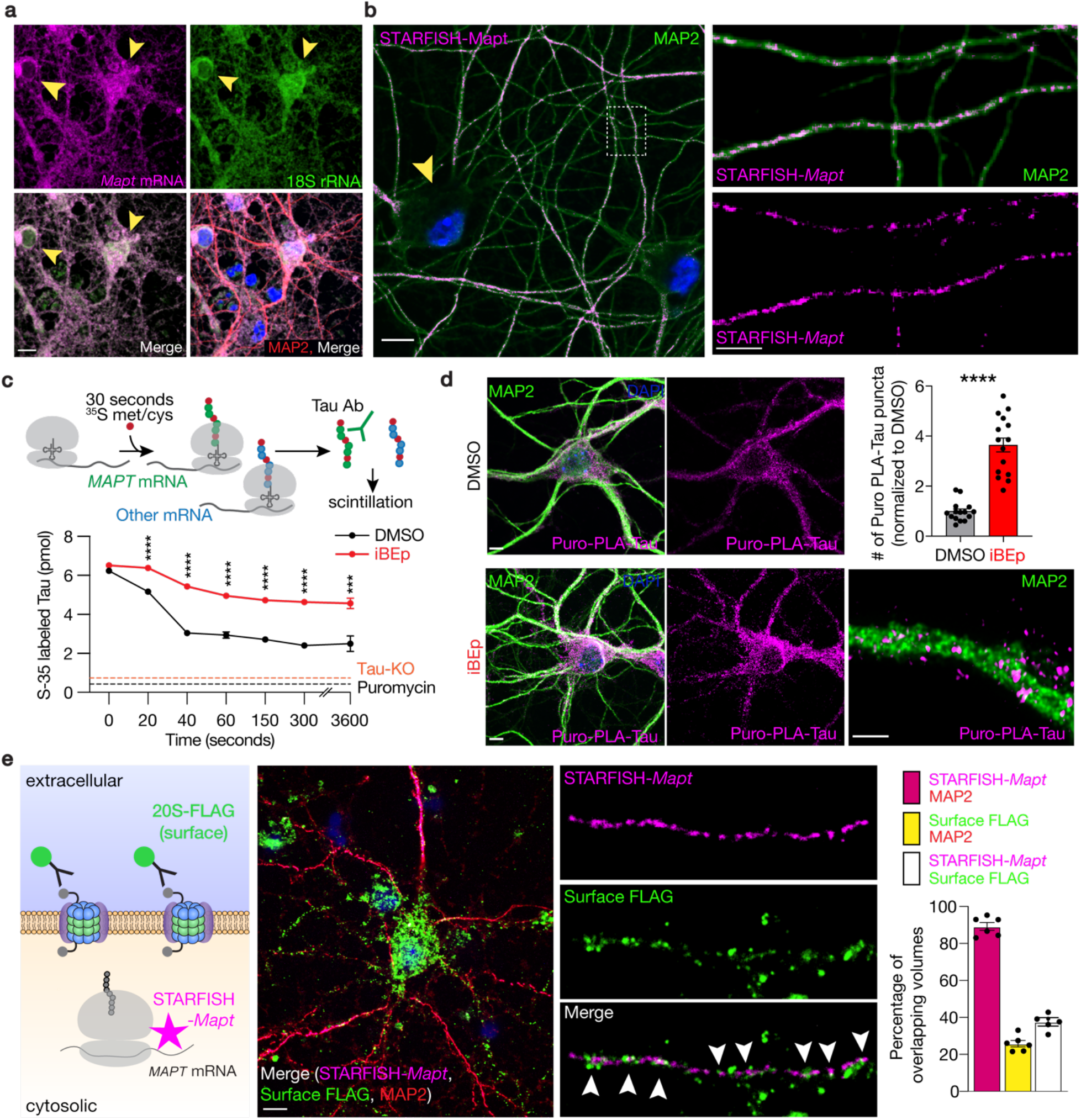
Tau is synthesized in neuronal dendrites and rapidly degraded by neuroproteasomes. **a**, smFISH experiment against *Mapt* in primary neurons. Alexa Fluor 647 against Mapt (magenta), Alexa Fluor 488 against 18S rRNA (green), Map2 (red) Scale bar= 5µm. **b**, STARFISH against *Mapt* mRNA (encoding Tau). STARFISH-*Mapt* signal (Magenta), MAP2 (Green), DAPI (Blue). Large insets (right) to exemplify STARFISH-*Mapt* signal relative to MAP2+ dendrite. **c**, Pulse-chase experiment to analyze synthesis and degradation kinetics of Tau. 35S met/cys radiolabel (red circle) pulse incorporated for 30 seconds onto DIV14 primary cortical neurons from WT animals pre-treated with either DMSO or iBEp (30 minutes). Tau immunoprecipitates after indicated chase times were quantified by liquid scintillation, total amount of Tau quantified based on specific activity of isotope. Identical experiments performed after concomitant 35S label with puromycin pulse (dashed black line) or from Tau KO mice (dashed red line). N=3 biological replicates. **d**, Micrographs of Puro-PLA-Tau signal following iBEp treatment. Primary DIV14 hippocampal neurons from WT mice treated with DMSO or iBEp (red) and pulse puromycylated. Puro-PLA-Tau labeling (pink), MAP2 (green), DAPI (blue). Quantification of number of Tau-PLA-Puro puncta normalized to DMSO. N=3 biological replicates. ****p<0.0001 by Paired T-test, Scale bar= 5 µm. **e**, STARFISH-*Mapt* labeling together with FLAG antibody feeding against endogenously tagged 20S proteasome to label neuroproteasomes. Illustrative schematic to left, magenta indicates STARFISH-*Mapt* signal, green indicates neuroproteasome signal. White arrows indicate co-localization between STARFISH-*Mapt* and neuroproteasome signal. Scale bar= 10 µm. Object-based co-localization analysis within 0.2 µm of STARFISH-Tau, Neuroproteasome, and MAP2+ dendrite signals. N=6 independent cultures.

The conventional method used to visualize newly synthesized proteins is Puro-PLA^6,37^. In this experiment, a positive proximity ligation amplification (PLA) signal between an antibody against puromycin and an antibody against the target of interest indicates newly synthesized protein. The dendritic localization of Tau translation was particularly surprising since other groups have reported that Puro-PLA experiments can visualize newly translated Tau protein in neuronal somata. We reproduce this somatic Puro-PLA-Tau signal, similar to published work (**Extended Data Fig. 4e**). We suggest the discrepancy between Puro-PLA-Tau and STARFISH-*Mapt* arises from the diffusion or transport of the puromycylated nascent polypeptide into soma (diffusion estimated to be around 50-70*μ*m in 3D space over 10 minutes)^18,19,38^. Because STARFISH is performed without puromycylation or without reliance on any diffusible or mobile signal, and has high spatial resolution, we are able to visualize the dendritic translation of Tau.

### Neuroproteasomes degrade newly synthesized Tau in dendrites

While the dendritic translation of Tau is unexpected, it is unclear whether this localization is related to the propensity of Tau to aggregate and mislocalize to the somatodendritic compartment in AD. We recently found that endogenous PHF Tau aggregates form and hyperphosphorylated Tau mislocalizes to the somatodendritic compartment upon failure of the neuroproteasome (Paradise, Konrad-Vicario et al 2025, in revision). Neuroproteasome inhibition could drive Tau aggregation directly by impairing Tau degradation, or indirectly, by perturbing another cellular factor involved in Tau proteostasis. We hypothesized that newly synthesized Tau could be degraded by neuroproteasomes based on three lines of evidence: 1) neuroproteasomes degrade newly synthesized proteins, 2) neuroproteasomes and Tau translation are located dendritically, and 3) neuroproteasome inhibition results in Tau aggregation.

To test this, we rapidly labeled newly synthesized proteins and immunoprecipitated Tau to determine its synthesis and degradation kinetics (**Fig. 2c**). We labeled newly synthesized proteins in primary mouse neurons with a 30-second pulse of ^35^S methionine/cysteine. The addition of puromycin, which induces dissociation of the ribosome from the mRNA^20^, during the labeling window reduced radiolabel in the Tau immunoprecipitates (**Fig. 2c, dashed black line**) to levels comparable to performing the IP from Tau KO neurons (**Fig. 2c, dashed red line**). Based on these important controls, we conclude that radiolabel in the Tau immunoprecipitation after 30 second ^35^S Met/Cys incorporation represents newly synthesized Tau. Remarkably, we find that nearly 50% of newly synthesized Tau is degraded within 30-60 seconds, which is largely prevented by the application of the neuroproteasome-specific inhibitor iBEp (Paradise, Konrad-Vicario et al 2025, in revision) (**Fig. 2c**). Epoxomicin mimics the effects of iBEp, with no further rescue of degradation (**Extended Data Fig. 5a**). Given there was no change in total scintillation signal in cell lysates or in Tau synthesis (**Extended Data Fig. 5b**), we conclude that iBEp slows Tau degradation but does not increase Tau translation or global translation. In further support of that conclusion, there was no change in translation or degradation of MAP2, another microtubule-associated protein, in identical experiments (**Extended Data Fig. 5c**). Finally, we find that after the rapid degradation phase, the remaining Tau protein remains stable for hours, consistent with the reported long half-life of Tau protein (**Fig. 2c**). We conclude that neuroproteasomes, and not cytosolic proteasomes, co- or peri-translationally degrade newly synthesized Tau.

Consistent with this, we observe a striking ∼3.5-fold increase in Puro-PLA-Tau signal in iBEp-treated neurons relative to DMSO controls (**Fig. 2d**). We observe no signal in neurons not treated with puromycin, in neurons where we block protein synthesis using Cycloheximide or Anisomycin during puromycylation (**Extended Data Fig. 5d**), or in Tau KO neurons (**Extended Data Fig. 5e**). We do not find an increase in the number of Puro-PLA-Actin (**Extended Data Fig. 5f-h**) or Puro-PLA-Tubulin (**Extended Data Fig. 5i-k**) puncta after iBEp treatment. We therefore suggest the increase we observe in Puro-PLA-Tau is not due to increased global translation, a conclusion supported by global quantitative proteomics following iBEp treatment (Paradise, Konrad-Vicario et al 2025, in revision). Rather, we suggest that the increase in Puro-PLA-Tau following iBEp treatment is due to impaired degradation of newly synthesized Tau.

For Tau to be co- or peri-translationally degraded by neuroproteasomes, ribosomes and *Mapt* mRNA must be adjacent to neuroproteasomes. To determine this, we sparsely labeled ribosomes in primary mouse hippocampal neurons from 20S-FLAG mice (Paradise, Konrad-Vicario et al in revision) using a low viral titer to express eGFP-L10a and performed FLAG antibody feeding to label neuroproteasomes (**Extended Data Fig. 5l**). We observe 27.3 ± 8.6% of ribosomes within 0.2 µm to neuroproteasome signal, of which more than 90% are in dendrites (**Extended Data Fig. 5l**). To image ribosomes translating *Mapt* mRNA adjacent to neuroproteasomes, we performed STARFISH against Tau while measuring neuroproteasome localization using FLAG antibody feeding. We find 37.6 ± 4.0% of STARFISH-*Mapt* signal within 0.2 µm of neuroproteasomes, indicating that neuroproteasomes are positioned to degrade newly synthesized Tau (**Fig. 2e**).

### *De novo* protein synthesis is required for neuroproteasome-dependent Tau aggregation

Given that newly synthesized Tau is degraded by neuroproteasomes and that neuroproteasome inhibition induces Tau aggregation, we hypothesized that translation would be required for neuroproteasome-dependent formation of Tau aggregates. To test this, we leveraged both iBEp and a mechanistically distinct neuroproteasome inhibitor Sulfo-MG (Paradise, Konrad-Vicario et al 2025, in revision). We incubated primary mouse neuronal cultures with either iBEp or Epoxomicin, or Sulfo-MG and its parent compound MG132, and then co-incubated these cultures with cycloheximide, which blocks translation elongation^39^. Cycloheximide blocked iBEp- and SulfoMG-induced Tau aggregation in hTau-KI primary neurons (**Fig. 3a,b**). We also tested whether inhibition of translation blocks endogenous Tau aggregation *in vivo* in hTau-KI mice. We first confirmed cycloheximide efficiently blocked protein synthesis *in vivo* without affecting cell health (**Extended Data Fig. 6a-d**), consistent with previous reports^40,41^. We stereotactically co-injected cycloheximide with iBEp contralateral to iBEp alone into the hippocampus. We find a significant decrease in sarkosyl-insoluble Tau in the hippocampus from mice co-injected with iBEp and cycloheximide compared to those injected with iBEp alone (**Extended Data Fig. 6e**,**f**). Additionally, we find that co-injection of cycloheximide with iBEp completely eliminated AT8+ staining compared to contralateral injection of iBEp alone (**Fig. 3c**). We also observe a complete return to baseline of ThioS+ staining in hippocampi co-injected with iBEp and cycloheximide relative to those injected with iBEp alone (**Fig. 3d**). Taken together, we conclude that neuroproteasomes co-translationally degrade newly synthesized Tau and a failure of this degradation results in endogenous Tau phosphorylation and aggregation *in vivo*.

**Figure 3:**
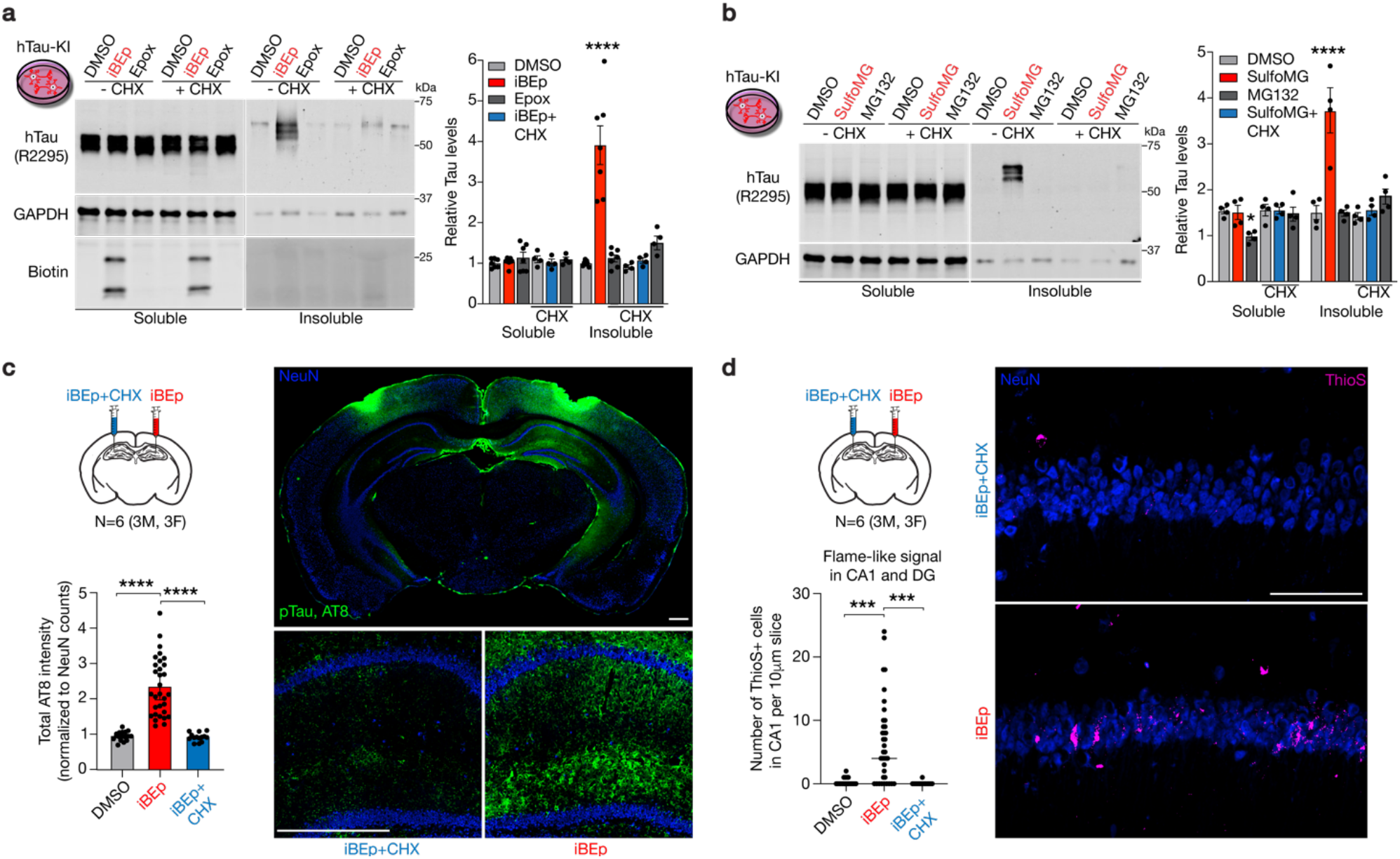
Translation is required for neuroproteasome inhibition-dependent Tau aggregation. **a**,**b**, Sarkosyl fractionation of primary hTau-knock-in (hTau-KI) neurons treated with (**a**) iBEp or Epox or (**b**) SulfoMG and MG132, concomitant with translation elongation inhibitor Cycloheximide (CHX). Sarkosyl-soluble and -insoluble fractions immunoblotted using indicated antibodies. LiCor-based quantification of relative Tau levels calculated as follows: Soluble and insoluble Tau intensities were normalized to respective soluble GAPDH intensity. All Tau levels plotted relative to DMSO in either soluble or insoluble fraction. Data (right) are mean ± SEM, N=4 independent biological replicates, iBEp + CHX or SulfoMG + CHX (blue) for emphasis relative to iBEp or SulfoMG alone (red). **c**, Immunohistochemical analysis of mice stereotactically injected with iBEp together with CHX to measure Tau phosphorylation *in vivo*. hTau-KI mice stereotactically injected into CA1 of hippocampus with iBEp ipsilaterally and coinjected with iBEp and CHX contralaterally. Mice were collected 72 hours post injection and sections were stained using indicated antibodies: NeuN (blue) and AT8 (pTau, green). Quantification of pTau signal intensity normalized to NeuN counts. Analysis was done blinded to experimental condition. Data are mean ± SEM normalized either to DMSO. N=6 (3M, 3F) biological replicates, n=3 sections/animal. Scale bar=500 µm. **d**, Thioflavin-S staining (to mark b-sheet containing aggregates) of sections from mice stereotactically injected with iBEp or iBEp with CHX. Conditions identical to (c), but stained with ThioS and NeuN. Data are quantification of number of flame-like Thioflavin-S positive inclusions. Counting and analysis was blinded to experimental condition. Scale bar=100 µm (right). ***p<0.001, ****p<0.001 by One-Way ANOVA Tukey’s Multiple Comparison Test. DMSO and iBEp data in (c,d) combined with experiments from Paradise, Konrad-Vicario et al, In revision

## Discussion

In this work, we present the development of a powerful tool to visualize the precise localization of new protein synthesis. The near-codon resolution afforded by STARFISH can visualize where within a cell or across cells that mRNA isoforms are translated or how 3’ or 5’UTRs modify the localization of translation. Beyond translational control, STARFISH would allow detection of specific deficits in protein translation, like stalling or skipping, or how disease-linked mutations affect translational output and localization. This is likely to be crucial in many disorders, as diverse disease states are characterized by dysfunctional and/or mislocalized ribonucleoproteins and transcripts. Beyond the applications in neurobiology and translational control, we envision applications for STARFISH and STARFISH derivatives in visualizing RNA binding protein interactions, impacts of RNA structure on localization and translation, and any instance where near-codon spatial resolution of RNA-protein interaction needs to be visualized.

Tau mislocalizes to the somatodendritic compartment in neurodegenerative diseases^12^. The prevailing rationale for this observation is that hyperphosphorylated Tau dissociates from microtubules in axons and diffuses into the somatodendritic compartment^5,10^. We suggest an additional explanation for this: Tau synthesis normally occurs in dendrites with ∼30% regularly degraded by neuroproteasomes and that neuroproteasome failure leads to Tau aggregation and phosphorylation in the somatodendritic compartment. It is counter-intuitive that Tau is synthesized in dendrites, far from where the majority of the folded protein is in the axon. We speculate this spatial segregation enables Tau synthesis and proteostasis in a protected compartment where potential misfolding events do not propagate in a compartment with a higher local concentration of Tau protein to seed further aggregation. Dendrites also have more abundant machinery for proteostasis than axons, potentially enabling neurons to handle the misfolding events in dendrites before misfolded Tau can reach the axon and spread. While we do not yet understand how Tau synthesis is restricted to dendrites, despite mRNA localization in the soma, it will be interesting to investigate the mechanisms of translational regulation in the future.

A striking finding from our work is that ∼30% of newly synthesized Tau is rapidly degraded by neuroproteasomes in healthy neurons, suggesting that neuroproteasomes are constitutively suppressing a baseline ‘overproduction’ of Tau by acting on the newly synthesized pool. When only neuroproteasomes are inhibited, that fraction of Tau is no longer degraded, and instead aggregates. This may appear to be energetically inefficient – however, we speculate two non-mutually exclusive hypotheses for future investigation to explain why this phenomenon occurs. The first is that degradation of Tau by neuroproteasomes releases Tau fragments into the extracellular space that have biological function. The second is that the newly synthesized pool of Tau that is targeted by neuroproteasomes is somehow defective^42^ and that neuroproteasomes degrade this pool as a quality control mechanism. In both cases, degradation of Tau by neuroproteasomes should result in secreted peptides derived from Tau in the extracellular space. In support of such proteolytic processing, N-terminal peptides from newly synthesized Tau were found to be secreted into CSF of human patients and in media of hiPSC-derived neurons^43^, albeit over days due to technical limitations of labelling Tau. Whether this phenomenon is neuroproteasome-dependent or has biological function is a subject of future study that will be aided by our development of neuroproteasome-specific inhibitors.

Exactly how Tau phosphorylation fits into our overall model is not yet clear. Following neuroproteasome inhibition *in vivo*, not all cells which are AT8 positive are ThioS positive – therefore, Tau phosphorylation (at least at the S202/T205 position) is not deterministic for aggregation under neuroproteasome-inhibited conditions. However, protein synthesis is required for both AT8 and ThioS positivity and for sarkosyl-insoluble Tau, following neuroproteasome inhibition. The AT8 positivity is therefore also not likely due to general cell stress induced by neuroproteasome inhibition - one might expect more stress with cycloheximide/iBEp coadministration or with Epoxomicin-based inhibition. Genetic and molecular details of the connection between neuroproteasomes and Tau will be essential to understand the overall relationships between Tau synthesis, degradation, phosphorylation, and aggregation. These will likely reveal intervention points that will be efficacious in delaying the onset and severity of Tau pathology in AD.

## Acknowledgements

We thank Ulrich Hengst, Clarissa Waites, and Ya-Cheng Liao for careful reading and comments, and Sadie for invaluable laboratory support. We thank Chi Nguyen for synthesis of iBEp and SulfoMG, David Holtzman for anti-Tau antibodies and protocol for Tau immunoprecipitation, David Schechner for help with designing DNA paint probes, and Rijuta Mukim for sections and immunostaining. We also thank Michael Shelanski, Richard Mayeux, and the Taub Institute for support and start-up funding.

## Author Contributions

KVR conceptualized the overall project while KDKV and KVR conceptualized the methodology for STARFISH together. KVR, VP, and KDKV designed methodology for all remaining experiments. KDKV performed nearly all experiments, except for Figure 3 which was performed by VP. LD conducted Schaffer collateral stimulations with equipment in CDM’s laboratory. KDKV helped with writing and KVR wrote the majority of the manuscript. KDKV and KVR assembled figures.

