## Supplemental Figures for "Spatial coupling of endogenous Tau translation and degradation by neuroproteasomes in dendrites revealed by STARFISH"

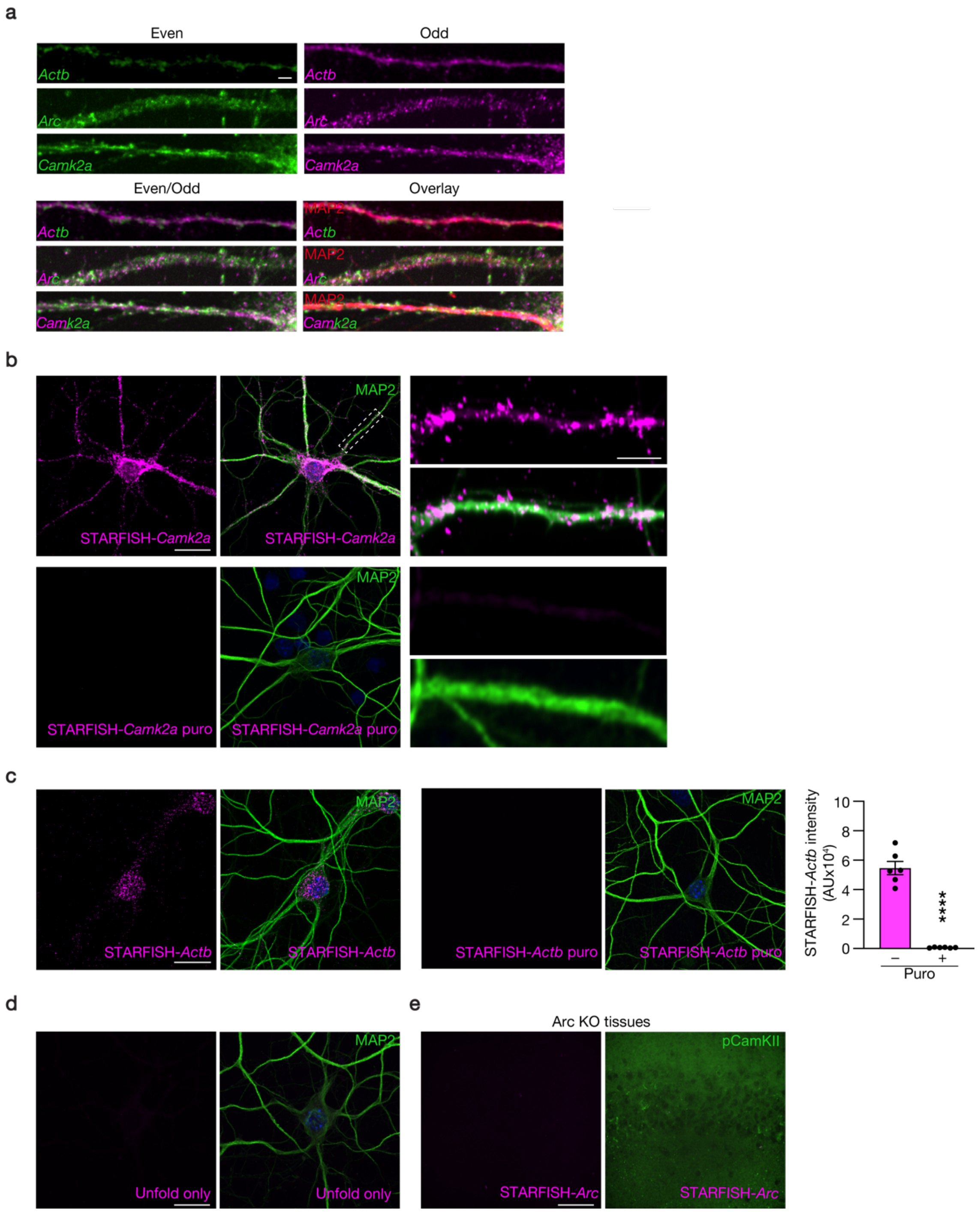


**Extended Data Figure 1. STARFISH measures the subcellular localization of translation with single-molecule sensitivity and high signal-to-noise**

**a,** Micrographs of smFISH from DIV18 primary hippocampal neurons with every other probe labeled in with Alexa Fluor 488 (even probes) or Alexa Fluor 647 (odd probes), MAP2 (red) Scale bar= 5µm. **b,** STARFISH against *Camk2a* mRNA (encoding CaMKIIa) in DIV18 primary hippocampal neurons. (top) STARFISH-*Camk2a* signal (Magenta), MAP2 (Green), DAPI (Blue), (bottom) STARFISH-*Camk2a* signal after Puromycin pulse. Insets to right of images, scale bar = 5 µm. **c,** STARFISH against *Actb* mRNA (encoding Beta-Actin) in DIV18 primary hippocampal neurons. (left) STARFISH-*Actb* signal (Magenta), MAP2 (Green), DAPI (Blue), (right) STARFISH-*Actb* signal after Puromycin pulse. N=6 biological replicates. Quantification of STARFISH signal intensity was normalized to total MAP2 area. Data are mean ± SEM. ****$p < 0.0001$ by Paired T-test. **d,** Unfold probes alone and Alexa Fluor rolling circle amplification performed without clickable mRNA or rRNA probes. **e,** STARFISH-*Arc* performed on slices from Arc KO animals. Unless stated otherwise Scale bar= 20µm.

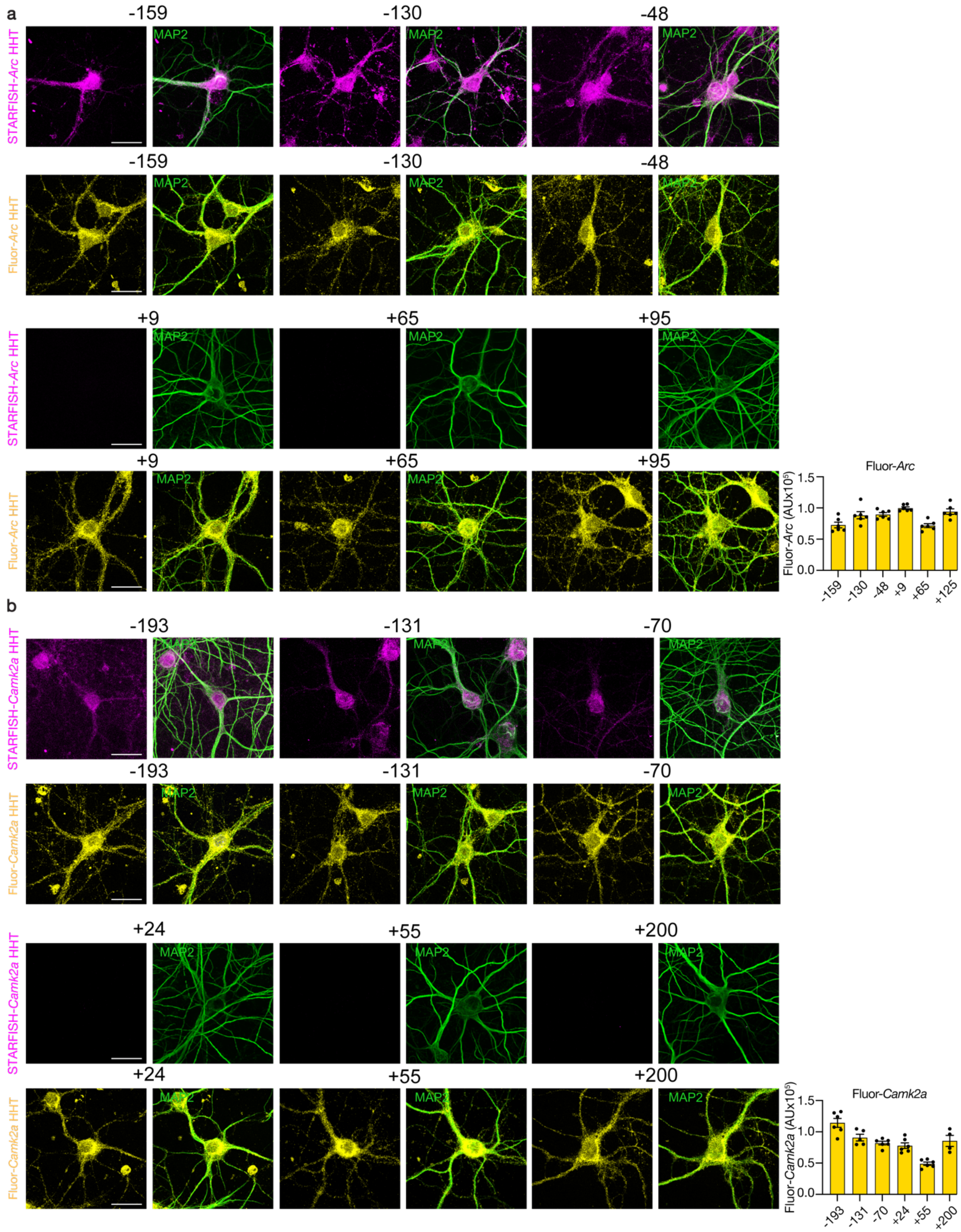


**Extended Data Figure 2. STARFISH measures the subcellular localization of translation with single-molecule sensitivity and near-codon resolution**

Homoharringtonine runoff experiments. **a,** STARFISH-*Arc* (Magenta) compared to Fluor-conjugated *Arc* mRNA probes (Yellow), Map2 (green). Single mRNA probes used for all experiments, mRNA probe position relative to the initiating AUG indicated. N=6 biological replicates. **b,** Same as (a) but with STARFISH-*Camk2a* (Magenta) and Fluor-conjugated *Camk2a* mRNA probes. Map2 (green). N=6 biological replicates. All scale bars= 20µm. Quantification of STARFISH signal or click fluorescence intensity was normalized to total MAP2 area. Data are mean ± SEM. ****p < 0.0001 by Paired T-test.

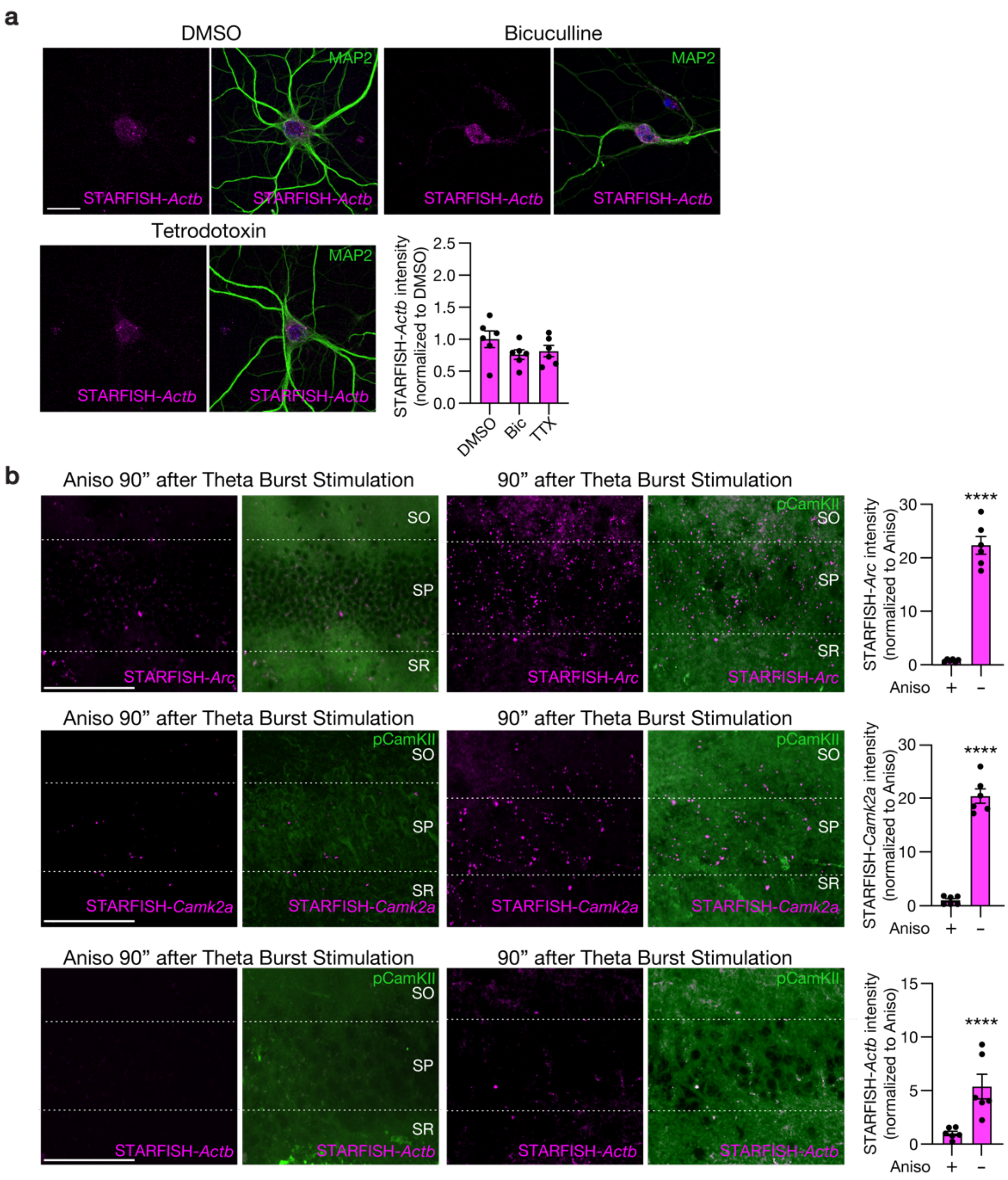


**Extended Data Figure 3. STARFISH detects activity-dependent translation in primary neurons, hippocampal slices, and *in vivo***

a, STARFISH against *Actb* mRNA in DIV18 primary hippocampal neurons. STARFISH-*Actb* signal (Magenta), MAP2 (Green), DAPI (Blue). STARFISH-*Actb* following stimulation of neuronal activity with Bicuculline or suppression with Tetrodotoxin. Quantification of normalized STARFISH intensity signals relative to DMSO. N=6 biological replicates. Scale bar=20 µm. b, STARFISH against *Arc*, *Camk2a*, and *Actb* in mouse hippocampal slices after theta burst stimulation. WT coronal hippocampal slices were either incubated in ASCF or ASCF with anisomycin prior to stimulation. Schaffer collaterals were stimulated in a burst pattern to induce robust activity in CA1 pyramidal neurons and fixed 90 seconds after stimulation. STARFISH was performed on tissue slices. N=2 biological replicates- 3 images taken of each

CA1 area. Scale bar=50 µm Quantification of STARFISH signal intensity was normalized to total MAP2 area. Data are mean ± SEM. ****$p < 0.0001$ by Paired T-test.

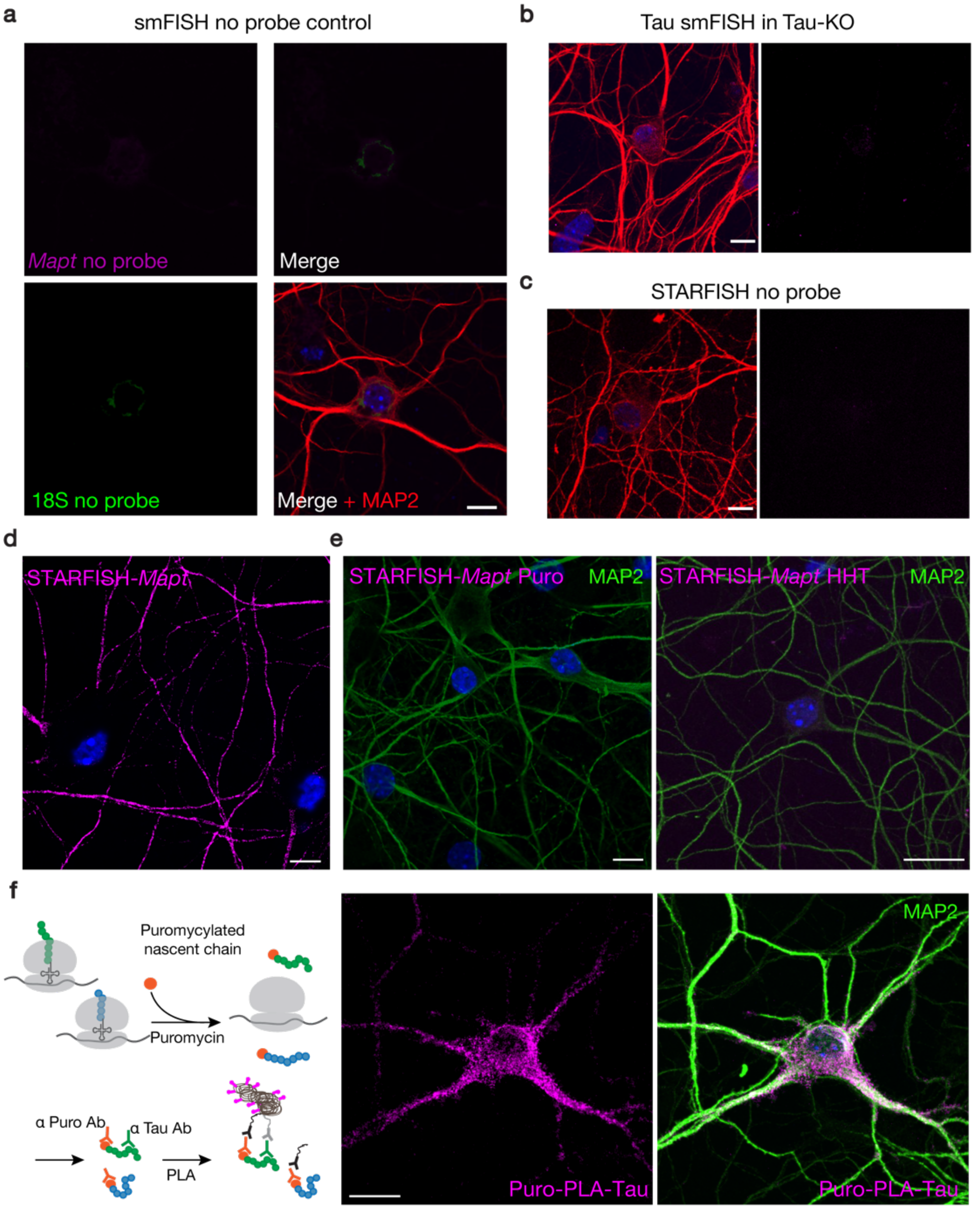


**Extended Data Figure 4. Tau is translated in neuronal dendrites**

**a,** Micrographs of smFISH from DIV18 primary hippocampal neurons using all secondary probes: Alexa Fluor 647 (magenta), Alexa Fluor 488 (green), Map2 (red) Scale bar= 5 µm. **b,** Micrographs of smFISH from DIV18 primary hippocampal neurons from Tau-KO mice using probes directed against *Mapt*(Tau) mRNA and 18S rRNA. 18S rRNA (green), *Mapt* mRNA (green), MAP2 (red). Scale bar= 5 µm. **c,** STARFISH assay against *Mapt* in primary neurons as in Fig 2b, but single channel image to emphasize presence of STARFISH-*Mapt* in majority of MAP2+ dendrites. **d,** STARFISH-Mapt in neurons treated with Puromycin (puro) or Homoharringtonine (HHT). Scale bare = 20 µm. **f,** Schematic of Puro-PLA (left), micrographs of Puro-PLA-Tau in primary hippocampal neurons. Scale bare = 5 µm.

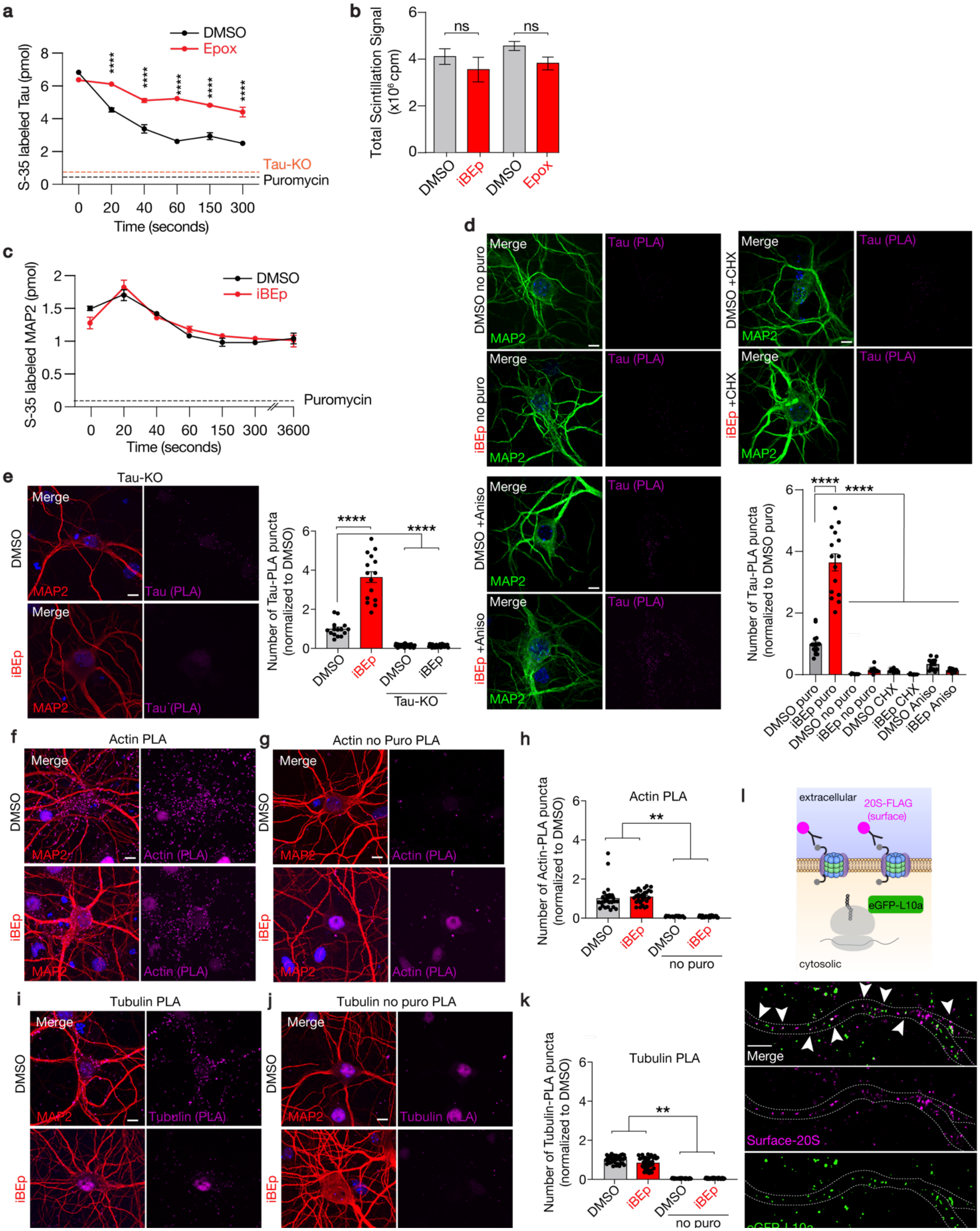


**Extended Data Figure 5. Newly synthesized Tau is degraded by neuroproteasomes**

**a,** Scintillation signal of Tau immunoprecipitates from DIV14 primary neurons pulse labeled with 35S met/cys for 30 seconds treated with indicated compounds and chased for indicated times. N=3 biological replicates, ****$p<0.0001$, ns = not significant by One-way ANOVA. **b,** Total scintillation signal of lysates from DIV14 primary neurons pulse labeled with 35S met/cys for 30 seconds treated with indicated compounds. N=3 biological replicates. ns = not significant by One-way ANOVA **c,** Scintillation signal of MAP2 immunoprecipitates from DIV14 primary neurons pulse labeled with 35S met/cys for 30 seconds treated with indicated compounds and chased for indicated times. Identical experiments performed after puromycin pulse concomitant with 35S pulse (dashed black line in graph). N=3 biological replicates, ns = not significant by One-way ANOVA. **d,** Micrographs of Puro-PLA-Tau signal in DIV14 primary WT hippocampal neurons from WT mice treated with DMSO or iBEp, in addition to either 1) not puromycylated (No Puro), 2) puromycylated in the presence of cycloheximide (CHX), or 3) puromycylated in the presence of anisomycin (Aniso). Tau-PLA-Puro labeling (pink), MAP2 (green), DAPI (blue). Quantification of number of Tau-PLA-Puro puncta were plotted together and normalized to DMSO Puro. ****$p<0.0001$ by One-Way ANOVA. Scale bar= 5 μm. **e,** Experiment as in Fig 2d, but in DIV14 primary hippocampal neurons derived from Tau KO animals. Quantification (right) of Tau-PLA-Puro signal was normalized to DMSO, data from Fig 2d plotted side by side for comparison to Tau KO. **f,** Micrographs of Actin-PLA-Puro labeling in DIV14 primary hippocampal neurons treated with iBEp. Actin-PLA-Puro (pink), MAP2 (green), DAPI (blue). Scale bar= 5 μm. **g,** Experiment as in (f) but not puromycylated. **h,** Quantification of Actin-PLA-Puro signal normalized to DMSO Puro. ****$p<0.0001$ by One-way ANOVA. **i,** Micrographs of Tubulin-PLA-Puro labeling in DIV14 primary hippocampal neurons treated with iBEp. Tubulin-PLA-Puro (pink), MAP2 (green), DAPI (blue). Scale bar= 5μm. **j,** Experiment as in (g) but not puromycylated. **k,** Quantification of Tubulin-PLA-Puro signal normalized to DMSO Puro. ****$p<0.0001$ by One-way ANOVA. N=3 biological replicates. **l,** Micrographs of primary neurons from 20S-FLAG mice expressing GFP-tagged ribosomes and surface labeled with anti-FLAG antibodies to label neuroproteasomes. Schematic above, micrographs below. White arrowheads indicate overlap.

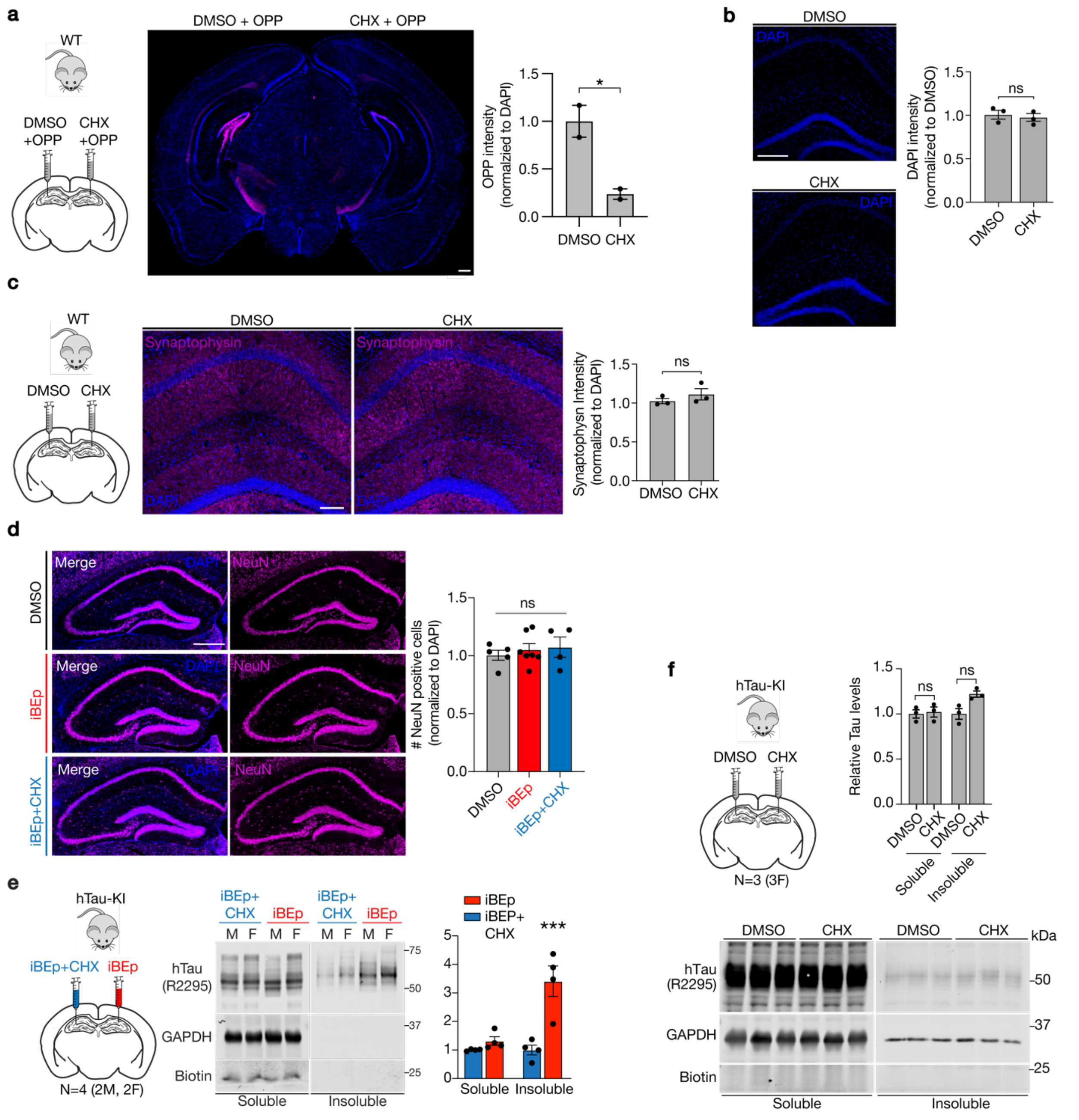


**Extended Data Figure 6: Translation is required for neuroproteasome inhibition-dependent Tau aggregation**

**a,** Immunohistochemical analysis of mice stereotactically injected with O-Propargyl Puromcyin (OPP) and OPP together with Cycloheximide. Janelia Fluor 646 was clicked onto OPP to visualize the site of puromycin incorporation. OPP signal quantified to right normalized to DAPI. N=2 biological replicates, 4 sections/animal. *p<.05 by Student T-Test. **b,** Micrographs from WT mice stereotactically injected with DMSO ipsilaterally and CHX contralaterally stained with DAPI. N=3 biological replicates, 4 sections/animal. DAPI signal quantified and plotted. ns = not significant by Student T-Test **c,** Micrographs of sections from WT mice stereotactically injected with indicated compounds. Sections were stained with Synaptophysin and DAPI.

Quantification of Synaptophysin intensity plotted normalized to DAPI intensity. N=3 biological replicates, 4 sections/animal. ns = not significant by Student T-Test **d,** Micrographs of sections from WT mice stereotactically injected with indicated compounds. Sections were stained with DAPI. Quantification of cell number by DAPI plotted. N=3 biological replicates, 4 sections/animal. ns by One-Way ANOVA. **e,** Immunoblots of sarkosyl soluble and insoluble fractions from hippocampi from hTau-KI mice stereotactically injected with iBEp ipsilaterally and iBEp + CHX co-injected contralaterally. Quantification of soluble and insoluble Tau intensities normalized to soluble GAPDH. Data shown (right) are mean ± SEM normalized to DMSO. N=4 biological replicates (2M, 2F), ***$p<0.001$ by One-Way ANOVA Tukey's Multiple Comparison Test. **f,** Immunoblots of sarkosyl soluble and insoluble fractions from hippocampi from hTau-KI mice stereotactically injected with DMSO and CHX alone and processed as in (e). N=3 biological replicates.
